# *Candida albicans*: a comprehensive view of the proteome

**DOI:** 10.1101/2024.12.20.629377

**Authors:** Leticia Gomez-Artiguez, Samuel de la Cámara-Fuentes, Zhi Sun, María Luisa Hernáez, Ana Borrajo, Aída Pitarch, Gloria Molero, Lucía Monteoliva, Robert L. Moritz, Eric W. Deutsch, Concha Gil

## Abstract

We describe a new release of the *Candida albicans* PeptideAtlas proteomics spectral resource (build 2024-03), providing a sequence coverage of 79.5% at the canonical protein level, matched mass spectrometry spectra, and experimental evidence identifying 3382 and 536 phosphorylated serine and threonine sites with false localization rates of 1% and 5.3%, respectively. We provide a tutorial on how to use the PeptideAtlas and associated tools to access this information. The *C. albicans* PeptideAtlas summary web page provides “Build overview”, “PTM coverage”, “Experiment contribution”, and “Dataset contribution” information. The protein and peptide information can also be accessed via the *Candida* Genome Database via hyperlinks on each protein page. This allows users to peruse identified peptides, protein coverage, post-translational modifications (PTMs), and experiments identifying each protein. Given the value of understanding the PTM landscape in the sequence of each protein, a more detailed explanation of how to interpret and analyse PTM results is provided in the PeptideAtlas of this important pathogen.

*Candida albicans* PeptideAtlas web page: https://db.systemsbiology.net/sbeams/cgi/PeptideAtlas/buildDetails?atlas_build_id=578

## INTRODUCTION

*Candida albicans* is a dimorphic fungus that is a component of the commensal human microbiota. However, under certain conditions, it can cause invasive candidiasis, particularly in immunocompromised and critically ill patients.^1,2^ This is one of the primary nosocomial infections, especially among patients in intensive care units, posing significant public health risk.^3,4^ Consequently, the World Health Organization (WHO) has included this organism in its priority list of fungal pathogens.^5^

As a result, the search for diagnostic and prognostic biomarkers of candidiasis has increased, as have efforts to better understand clinically relevant biological processes for intervention. This has led to the development of multiple proteomics studies to understand the proteome complement and identify potential targets of intervention. However, despite the extensive efforts on clinical aspects from the proteomics view, there were no *C. albicans* online public proteomic repositories until the first *C. albicans* PeptideAtlas was published in 2014.^6^ PeptideAtlas is a multi-organism, publicly accessible compendium of mass spectrometry (MS) identified peptides identified from a large set of tandem mass spectrometry proteomics experiments through data reprocessing of MS datasets available through ProteomeXchange^7^ with the Trans-Proteomic Pipeline (TPP)^8^ and makes an integrated view of the results presented back to the community via a consolidated web resource. In spite of being heavily focused on the human proteome since the first publication in 2005^9^ PeptideAtlas has also created builds for multiple other species, such as pig (*Sus scrofa*),^10^ chicken (*Gallus gallus*),^11^ plant species (*Arabidopsis thaliana*^12^ and *Zea mays*^13^), several yeast species (*Saccharomyces cerevisiae,*^14^ *Schizosaccharomyces pombe,*^15^ and *C. albicans*^6^) and some bacteria species (*Staphylococcus aureus*^16^ and *Pseudomonas aeruginosa*^17^), all available under a common interface.

In the 2014 *C. albicans* PeptideAtlas, we provided a significant large-scale public proteomic resource for the study of this opportunistic pathogenic fungus. It initially cataloged over 2,500 proteins, covering approximately 41% of the predicted proteome. This achievement marked unprecedented proteome coverage for *C. albicans* and distinguished it as the first human fungal pathogen included in the PeptideAtlas project. However, this initial coverage was notably lower than that of other yeast species, such as *S. cerevisiae* (61%)^14^ and *S. pombe* (71%).^15^ Consequently, a subsequent update^18^ expanded the dataset to 4,115 identified proteins, thereby increasing the proteome coverage to 66% and bringing it closer to the levels observed in these other yeast species.

Since the last update in 2016, there has been a notable increase in the number of proteomic studies on *C. albicans* (*Supplementary Figure 1*), making it feasible to undertake a second update of the PeptideAtlas. A rigorous search was conducted on Proteomics Identifications Database (PRIDE),^19^ a proteomics repository, selecting studies that adhered to established criteria. These novel datasets, along with previously archived raw MS data files from the two preceding versions of the *C. albicans* PeptideAtlas, have undergone comprehensive reprocessing and analysis through the TPP^8,20^ utilizing an identified peptide sequence database featuring allele-specific sequences from the *Candida* Genome Database (CGD) in conjunction with one from UniProtKB. The outcome is the generation of the most comprehensive *C. albicans* proteomics resource to date, featuring unprecedented proteome coverage and incorporating, as a novelty, the calculation of the probabilities of two post-translational modifications (PTMs) in the amino acid sequence, phosphorylation and acetylation. Considering both the existing and newly incorporated tools, we present a tutorial for utilizing the *C. albicans* PeptideAtlas. Additionally, we provide updates on the process to develop the resource, and the improvements made.

## MATERIAL AND METHODS

### a) Selection of European Bioinformatics Institute (EBI) PRIDE submissions

ProteomeXchange Dataset selections (i.e., PXD records) were defined in PRIDE^19^ according to various criteria: submissions from 2015 onwards, studies involving the strain SC5314 widely used in the *C. albicans* field or its derived mutants, proteomic analysis conducted via LC-MS/MS and DDA, absence of protein glycosylation enrichment, and utilization of the CGD for peptide assignment. Thirty-three PXD datasets downloaded met these criteria and detailed information about them can be found in *Supplementary Table 1 (xlsx)*.

### b) The TPP data processing pipeline

For each PXD, we gathered data information regarding the protease, whether the samples were isotopically labeled, PTM enrichment techniques utilized, and the *Candida* strains selected for the experiments. According to the information collected, the runs of each PXD dataset were organized into one or more sub experiments. MS vendor-format raw files were converted to mzML files using ThermoRawFileParser^21^ for Thermo Fisher Scientific instruments, AB_SCIEX_MS_Converter for SCIEX wiff files, and tdf2mzml for Bruker timsTOF raw files. mzML files were searched using MSFragger,^22^ then processed through the TPP^8,20,24^ v6.4.x Pillar, Build 202403120129-9139 for peptide and protein identification. The protein search database included (i) Assembly 22 (A22-s07-m01-r202) (http://www.candidagenome.org/) with 6,226 sequences, (ii) UniProtKB reference proteome UP000000559^23^ with 6,036 sequences, (iii) 498 contaminant protein sequences frequently observed in proteome samples and (iv) a sequence-shuffled decoy counterpart, For the database searching, the following parameters were used: search_enzyme_name_1=trypsin, allowed_missed_cleavage_1=2, and num_enzyme_termini=1. The following variable modifications were set: oxidation of Met or Trp (+15.9949), peptide N-terminal Gln to pyro-Glu (−17.0265), peptide N-terminal Glu to pyro-Glu (−18.0106), deamidation of Asn (+0.9840), protein N-term acetylation (+42.0106), Ser, Thr, Tyr and Ala (+79.9663) if phosphorylated peptides were enriched, acetylation of Lys (+42.0106) if acetyl-peptides were enriched, hydroxyisobutyryl of Lys (+86.036779) if 2-hydroxyisobutyrylation peptides were enriched, and crotonyl of Lys (+68.026215) if lysine crotonylation enrichment was performed. Static carbamidomethylation of Cys (+57.0215) was set for all datasets except PXD003685 for which carbamidomethylation of Cys (+57.0215) and glutathione of Cys (+305.068156) were set as variable modifications. Falsely phosphorylated alanine was used as a decoy for phosphorylation localization analysis as described by Ramsbottom *et al.*^25^ For isotopically labeled samples with tandem mass tag (TMT) or SILAC, appropriate mass modifications were applied. PeptideProphet^26^ was used to assign peptide-spectrum match (PSM) probabilities on the sequence database search results. These probabilities were further refined using the iProphet tool.^27^ For acetylated and phosphorylated peptide enrichment experiments, PTMProphet^28^ was run to obtain localization probabilities.

### c) PeptideAtlas assembly

All datasets were thresholded at a probability that yields an iProphet model-based global false discovery rate (FDR) of 0.001 at the spectrum level. The exact probability threshold varied from experiment to experiment depending on how well the modeling can separate correct from incorrect information. iProphet probability 0.9 was used as the minimum even when global FDR analysis would favor a lower probability. Throughout the procedure, decoy identifications are retained and then used to compute final decoy-based FDRs. The decoy-based PSM-level global FDR is 0.00017, the peptide sequence-level global FDR is 0.0008, and the protein-level global FDR is 0.007 according to the MAYU^29^ analysis.

Proteins are identified using standardized assignments to different confidence levels based on various attributes and relationships to other proteins using a 10-tier system^12^ developed over many years for the human proteome PeptideAtlas^30,31^ (*Supplementary Table 1 (xlsx)*). The highest confidence level category is the “canonical” or “non-core canonical” category, which requires at least two uniquely mapping non-nested (one not inside the other) peptides at least 9 aa long with a total coverage of at least 18 aa, as required by the HPP guidelines,^32^ The decoy-based canonical protein FDR is 0.0005 with 2 decoys and 5,289 target canonical and noncore-canonical proteins including contaminants.

When a group of proteins cannot be disambiguated because they contain shared peptides, one or more “leaders” of the group are categorized as “indistinguishable representative” or “representative” *(Supplementary Table 2)*. This means that the protein or one of its close siblings is detected, but it is not possible to disambiguate them. The “marginally distinguished” category means that the protein shares peptides with a canonical entry but has some additional uniquely mapping peptide evidence that is however not sufficient to raise it to the canonical level. The “weak” category means that there is at least one uniquely mapping peptide that is nine or more residues long, but the evidence does not meet the criteria for being canonical. The “insufficient evidence” category means that all the uniquely mapping peptides are less than nine residues long. Finally, all other proteins that lack any matched peptides observed above our minimum PSM significance threshold are categorized as “not observed” proteins.

## RESULTS AND DISCUSSION

The 2024 update of the *C. albicans* PeptideAtlas includes several key steps (*Figure 1*). Initially, suitable datasets were selected from PRIDE based on specific criteria. The MS runs from the 33 new *C. albicans* datasets^33–53^ were then organized according to protease, labeling, enrichment technique, and *C. albicans* strain used. Subsequently, these new datasets, along with one archived dataset from the two preceding versions of the *C. albicans* PeptideAtlas, were reprocessed and analyzed using the TPP (version TPP v6.4.x Pillar, Build 202403120129-9139), a comprehensive end-to-end data analysis pipeline that comprises four distinct programs: PeptideProphet, iProphet, PTMProphet, and ProteinProphet. Finally, all the data was assembled taking into account the FDR employed in each program and the confidence level of the proteins identified.

**Figure 1.**
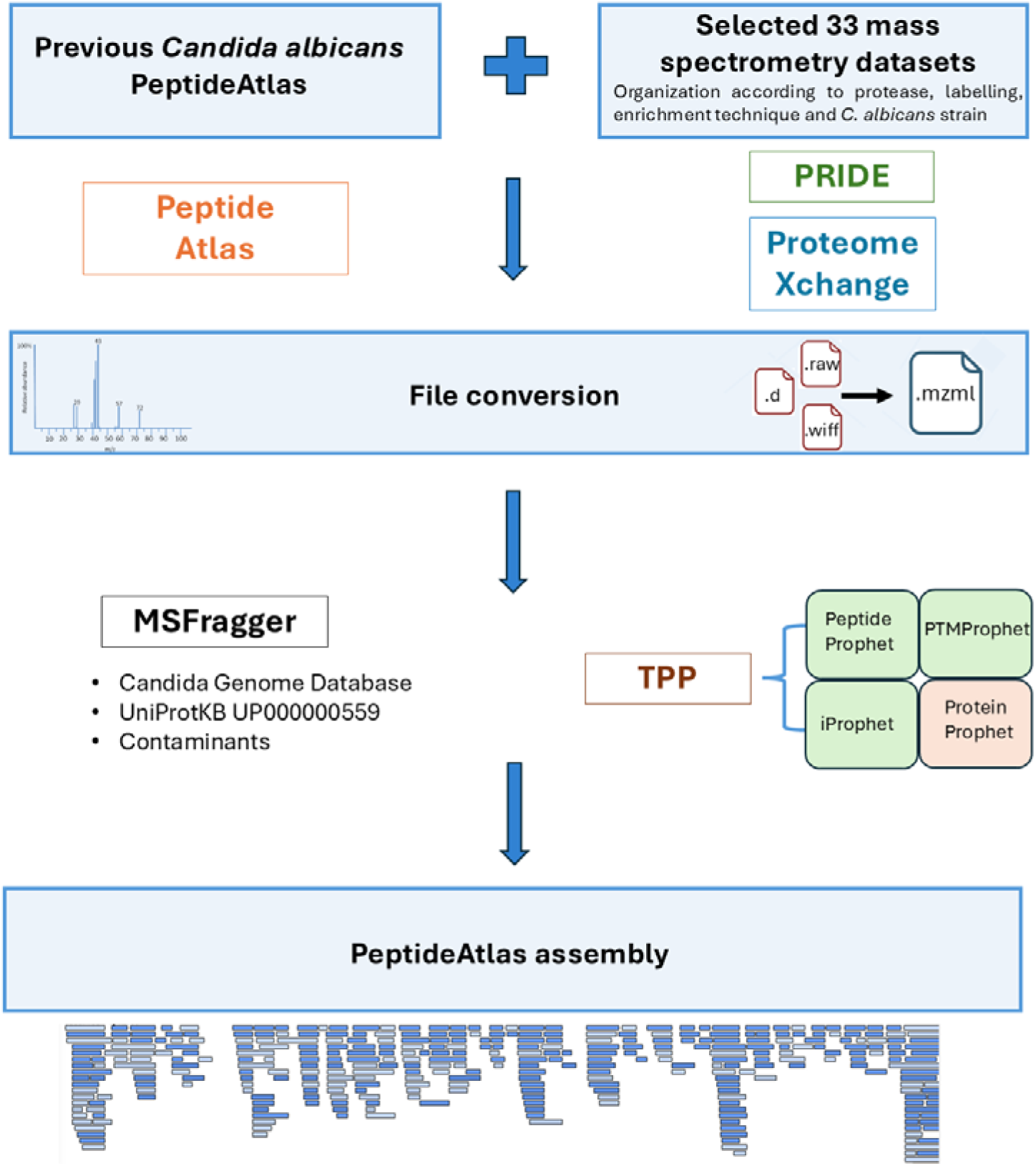
Graphical overview of the *C. albicans* PeptideAtlas upgrade. Specific steps are shown in blue boxes.

### a) *C. albicans* Protein coverage

A total of 834 MS runs (605 corresponding to the 33 datasets incorporated for the 2024 PeptideAtlas update, plus 148 from the 4 experiments added in the previous *C. albicans* PeptideAtlas build, plus 81 from the 16 experiments included in the first *C. albicans* PeptideAtlas build) generated 32,063,951 spectra of which more than one-third, 11,925,423, could be allocated to a peptide sequence. In the resulting outcome, for a decoy-based PSM FDR threshold set at 0.02%, 176,568 peptides are detected which can be explained by the minimal non-redundant set of 4,955 canonical *C. albicans* Assembly #22 protein sequences representing 79.5% of the 6,226 (as of May 2024) predicted different protein sequences.

The newly included LC-MS/MS datasets represent an increase of more than 12-fold in identified PSMs, 2.5-fold in terms of peptides and 1.2-fold in the number of identified canonical proteins. The improvements regarding the previous and current versions of the *C. albicans* PeptideAtlas are summarized in *Table 1*.

**Table 1.** Summary of the information included in the 3 PeptideAtlas performed to date (2011-12, 2015-02, 2024-03) and the improvements they have entailed.

| **PEPTIDEATLAS version** |  | **2011-12<sup>6</sup>** | **2015-02<sup>18</sup>** | **2024-03** |
| --- | --- | --- | --- | --- |
| Datasets | Datasets incorporated | 1 | 1 | 34 |
|  | Experiments analyzed | 16 | 20 | 113 |
| RUNS | Runs incorporated* | 81 | 148 | 605 |
|  | Runs analyzed ** | 81 | 229 | 834 |
| Spectra | New identified spectra | 172,941 | 811,142 | 10,941,340 |
|  | Identified spectra | 172,941 | 984,083 | 11,925,423 |
|  | Fold increase | - | 5.69 | 12.12 |
| Peptides | New distinct peptides | 21,938 | 49,372 | 105,258 |
|  | Distinct peptides identified | 21,938 | 71,310 | 176,568 |
|  | Fold increase | - | 3.25 | 2.48 |
| Proteins | New canonical proteins | 2,563 | 1,553 | 690 |
|  | Canonical proteins identified | 2,563 | 4,225 | **4,955** |
|  | Fold increase | - | 1.65 | 1.17 |
| Database | Version | A21-s02-m05-r10 | A22-s05-m01-r01 | A22-s07-m01-r202 |
|  | Nº proteins | 6,209 | 6,218 | 6,226 |
| Coverage | Percentage | 41.3% | 66.2% | 79.5% |
|  | Fold increase | - | 1.60 | 1.20 |

\* Number of runs included in previous versions.
\*\*Total number of runs analyzed.

One remarkable value provided by the *C. albicans* PeptideAtlas is the report of highly confident identification of proteins corresponding to uncharacterized genes (following the terminology in CGD), i.e. those genes without previously known empirical evidence for a translated product. These amounted to 2,860 in the previous PeptideAtlas build and have notably increased to 3,180, representing 76% of the total uncharacterized genes in CGD (*Figure 2*). As for the verified set of genes, those that do have experimental evidence for a gene product, 87% are covered in the list of canonical proteins in this build, increasing from 1,172 identified in the previous one to 1,623. Finally, as a novelty, one dubious gene, which is unlikely to encode a product and appears indistinguishable from random non-coding sequences, was identified as a canonical protein, representing 1% of the total dubious genes.

**Figure 2.**
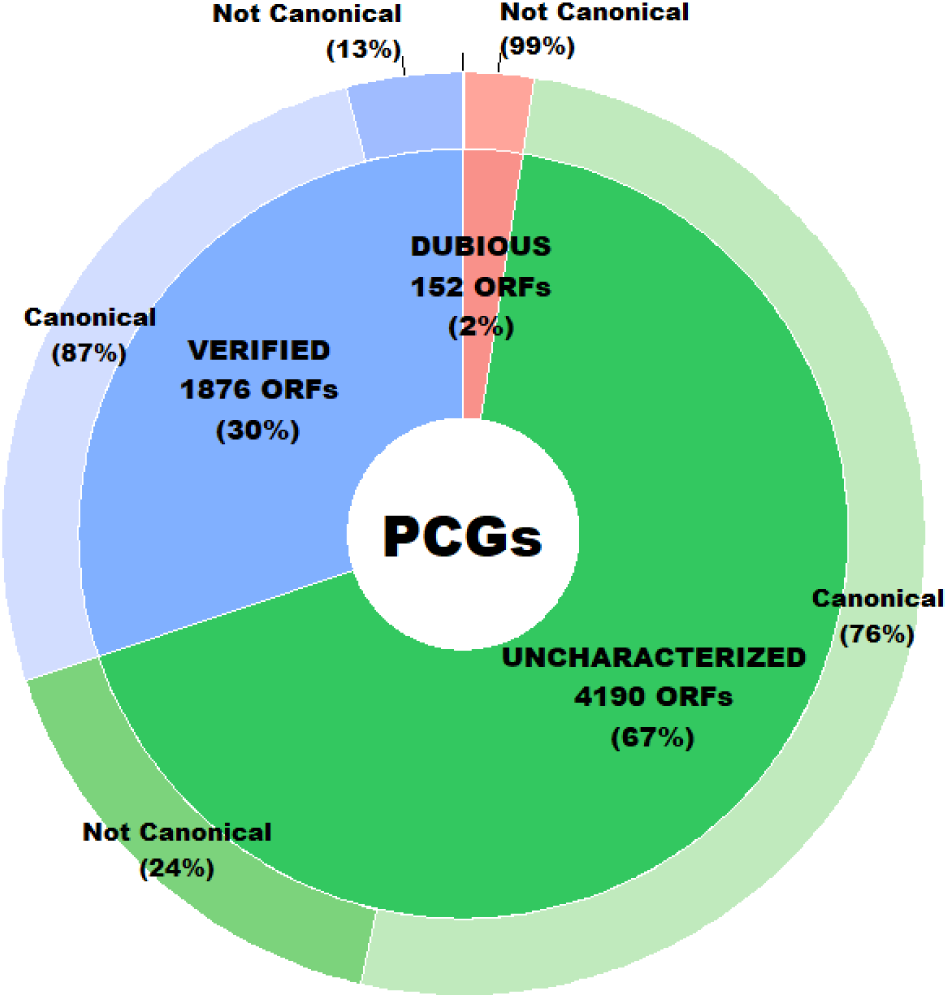
Classification of the 6,218 protein coding genes (PCGs) of *C. albicans* in Uncharacterized, Verified and Dubious and the percentage of these covered by canonical proteins (high confidence) identified in *C. albicans* PeptideAtlas.

An overview of the contributions of individual experiments to the entirety of the PeptideAtlas is depicted in *Figure 3*. *Figure 3A*, shows the cumulative identified peptides as well as distinct peptides from the 834 experiments whereas *Figure 3B* shows the cumulative identified canonical proteins as well as distinct canonical proteins from the 834 experiments. In both cases a certain region of the figure is marked for comparison in order to highlight the experiments that constituted the previous PeptideAtlas build. The build is based on 834 experiments across the 53 selected PXDs, where each PXD may be decomposed into several sub experiments. Areas in orange and red in *Figure 3* indicate the total number of distinct peptides or proteins respectively in each experiment, whereas areas in blue indicate the cumulative number from the current and previous experiments.

**Figure 3.**
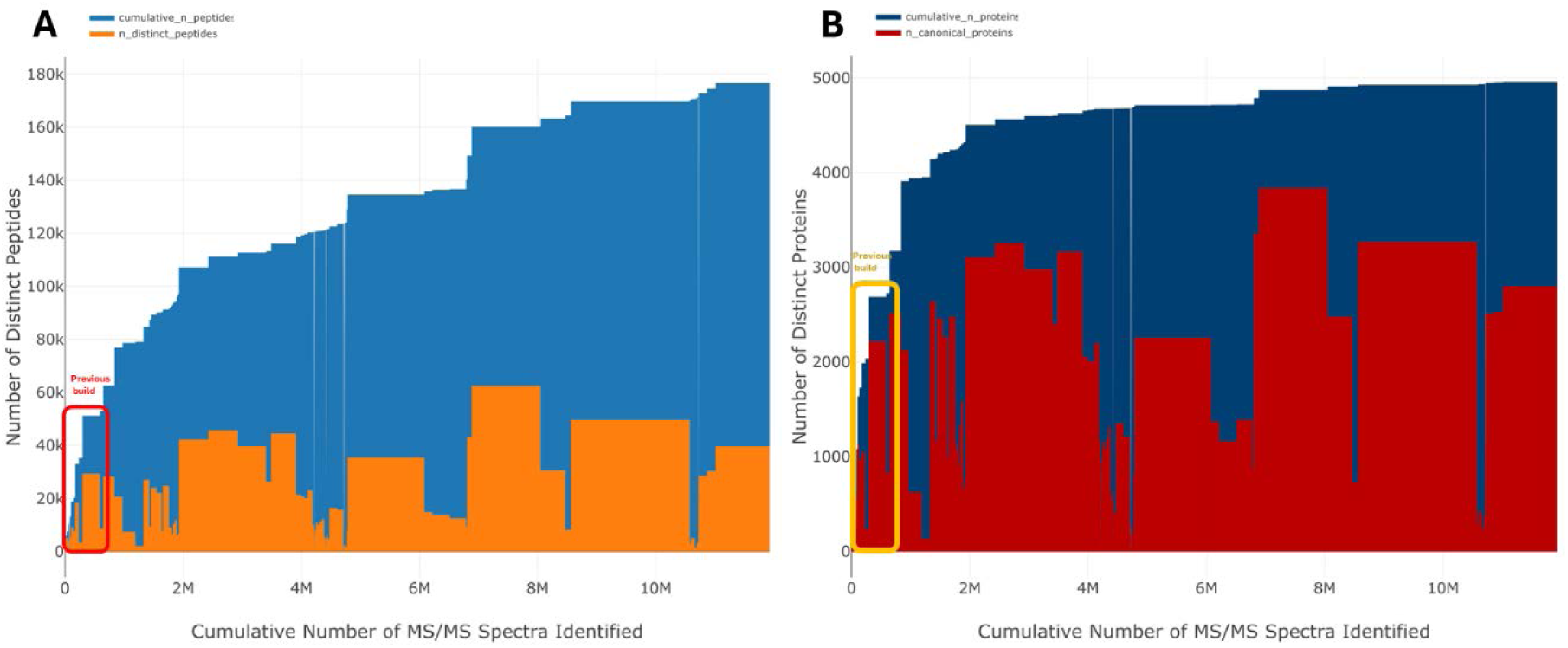
Number of distinct (non-redundant) peptides (A) and identified canonical proteins (B) as a function of the cumulative number of PSMs (peptide-spectrum matches) for the *C. albicans* PeptideAtlas update. The cumulative count is ordered by PXD identifier (from old to new).

For the better understanding of the underlying data of this PeptideAtlas build, we calculated the frequency distributions of peptide charge state and peptide length for the approximately 12 million matched MS/MS spectra (*Figure 4A*). The majority of matched spectra had a charge state of 2+ (57.19%) followed by 3+ (36.42%) and 4+ (5.78%). The charge states that yield the smallest number of spectra were 5+ and 1+ with 0.12% and 0.48% respectively. We observed a wide range of matched peptide lengths, with 7 aa being the shortest sequence allowed (*Figure 4B*). In addition, 98.8% of all matched peptides were between 7 and 37 aa long, with the most frequent peptide length of 12 aa (7.36%).

**Figure 4.**
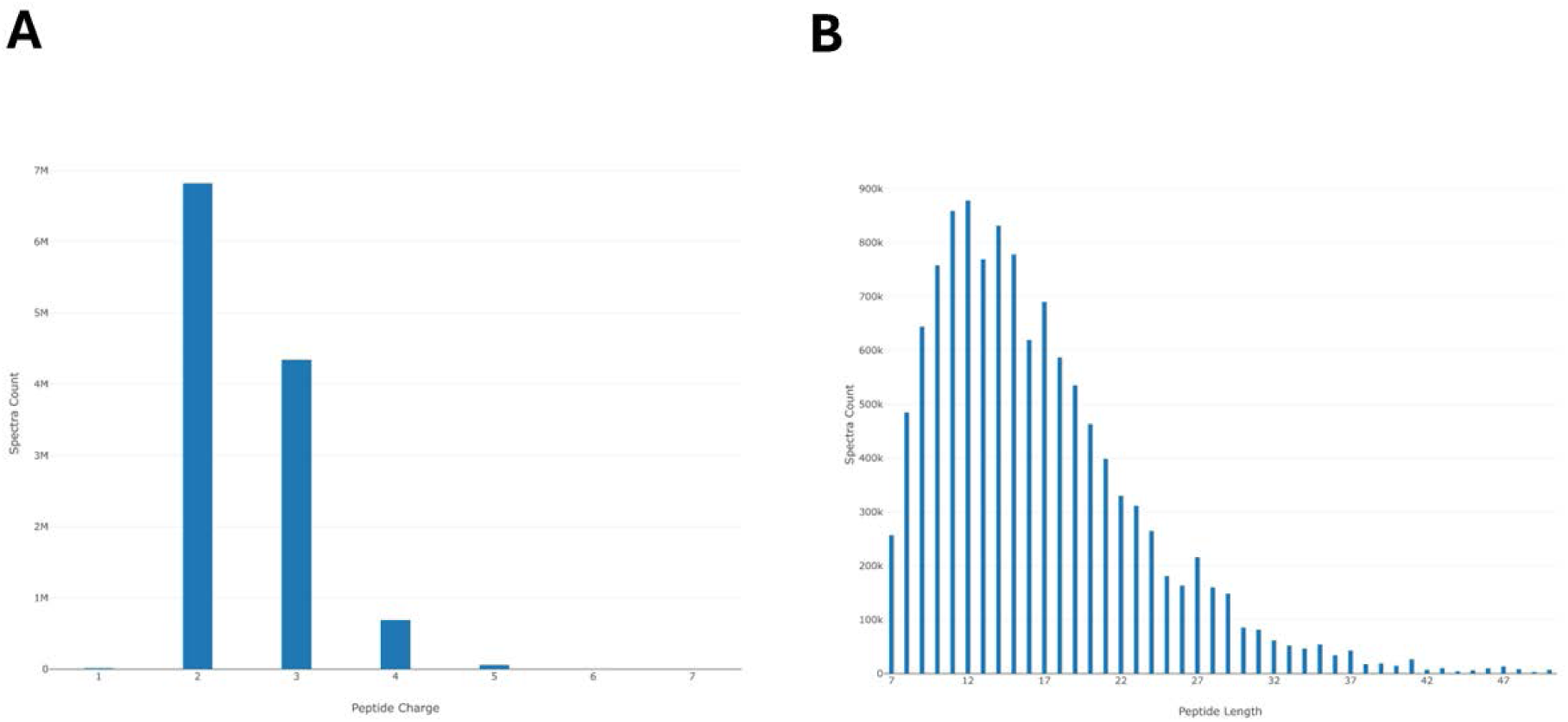
Key statistics of matched MS/MS data for *C. albicans* PeptideAtlas. (A) Frequency distribution of peptide charge state (z). (B) Frequency distribution of peptide length (aa).

At the highest level of confidence, 4,955 canonical proteins were identified in CGD (*Table 2*), appearing to be equally distributed along the nuclear chromosomes, with each chromosome having approximately 80% of its proteins identified. Additionally, 9 proteins were categorized as “Indistinguishable representative”, 16 as “Insufficient evidence”, 38 as “Marginally distinguished” and 239 as “Weak” (*Table 2*). Finally, 956 (956/6,213 = 15.4%) predicted proteins in CGD were not identified. These “Not observed” proteins were also fairly evenly distributed across the eight nuclear chromosomes. A proteome coverage of approximately 80% represents a reasonable proportion of identified proteins. However, the remaining 20% of unidentified proteins can be attributed to several factors. One significant reason is the presence of low abundance proteins which may be below the detection limit of the instrument, making their detection challenging, physico-chemical properties that do not allow their detection by the technique, proteins that are not expressed under the experimental conditions chosen or because of the nature of the primary protein sequence, in particular the absence of enzymatic cleavage sites.

**Table 2.** Proteins identified using CGD (A22-s07-m01-r202) were classified according to the six PeptideAtlas protein categories and their respective chromosomes, including the nuclear chromosomes (1–7, R) and the mitochondrial chromosome (M).

|  | Entries | Canonical | % | Indistinguishable<br>representative | % | Insufficient<br>evidence | % | Marginally<br>distinguished | % | Weak | % | Not<br>observed | % |
| --- | --- | --- | --- | --- | --- | --- | --- | --- | --- | --- | --- | --- | --- |
| **C1** | 1,383 | 1,144 | 82.7 | 3 | 0.2 | 4 | 0.3 | 3 | 0.2 | 47 | 3.4 | 182 | 13.2 |
| **C2** | 1,017 | 813 | 79.9 | 1 | 0.1 | 4 | 0.4 | 7 | 0.7 | 32 | 3.1 | 160 | 15.7 |
| **C3** | 760 | 588 | 77.4 | 1 | 0.1 | 2 | 0.3 | 6 | 0.8 | 44 | 5.8 | 119 | 15.7 |
| **C4** | 674 | 523 | 77.6 | 2 | 0.3 | 1 | 0.1 | 5 | 0.7 | 32 | 4.7 | 111 | 16.5 |
| **C5** | 523 | 416 | 79.5 | 0 | 0.0 | 2 | 0.4 | 3 | 0.6 | 20 | 3.8 | 82 | 15.7 |
| **C6** | 439 | 356 | 81.1 | 1 | 0.2 | 0 | 0.0 | 3 | 0.7 | 19 | 4.3 | 60 | 13.7 |
| **C7** | 407 | 323 | 79.4 | 0 | 0.0 | 0 | 0.0 | 4 | 1.0 | 11 | 2.7 | 69 | 17.0 |
| **CR** | 990 | 783 | 79.1 | 1 | 0.1 | 3 | 0.3 | 6 | 0.6 | 30 | 3.0 | 167 | 16.9 |
| **CM** | 20 | 9 | 45.0 | 0 | 0.0 | 0 | 0.0 | 1 | 5.0 | 4 | 20.0 | 6 | 30.0 |
| **TOTAL** | 6,213 | 4,955 | 79.8 | 9 | 0.1 | 16 | 0.3 | 38 | 0.6 | 239 | 3.8 | 956 | 15.4 |

To enhance proteome coverage, support for Data-Independent Acquisition (DIA) data in future versions of PeptideAtlas will be a strong point that would increase the coverage achieved. In addition, improvements in instrumentation, including the use of more sophisticated and highly sensitive mass spectrometers, will further contribute to expanding coverage. In parallel the use of different approaches to analyze specific subcellular fractions could improve the assessment of the proteome.

### b) PeptideAtlas tools

The *C. albicans* 2024-03 PeptideAtlas build is accessible through its web interface at https://db.systemsbiology.net/sbeams/cgi/PeptideAtlas/buildDetails?atlas_build_id=578. The general information page is displayed which includes a “Build Overview” section containing information about the number of datasets, experiments and MS runs, the reference database employed and the number of identified spectra, distinct peptides and proteins classified into different protein presence levels. A “PTM coverage” section shows the number of PTM sites for phosphorylation and acetylation at several levels of confidence. Finally, it is necessary to highlight the “Experiment Contribution” and “Datasets Contribution” sections that display the contribution of each experiment and dataset respectively for the actual *C. albicans* build in terms of spectra, peptides and protein.

Furthermore, a linkage between CGD and PeptideAtlas ensures easy accessibility. By searching for a protein of interest in the CGD and clicking on “View peptides from PeptideAtlas” in the “Protein” section, users can access all PeptideAtlas tools. This includes viewing identified peptides in the “Sequence Motifs” section, obtaining detailed information about them either by clicking on them in this section or in the “Distinct Observed Peptides” section, assessing protein coverage in the “Sequence” section, and analyzing the probability of PTMs such as phosphorylation and acetylation within the amino acid sequence in the “PTM Summary” section. Additionally, users can see which experiments identified the protein of interest in the “Experiment Peptide Map” section. This allows observation of the expression of the 40 most highly observed peptides of the searched protein across various experiments (*Figure 5*).

**Figure 5.**
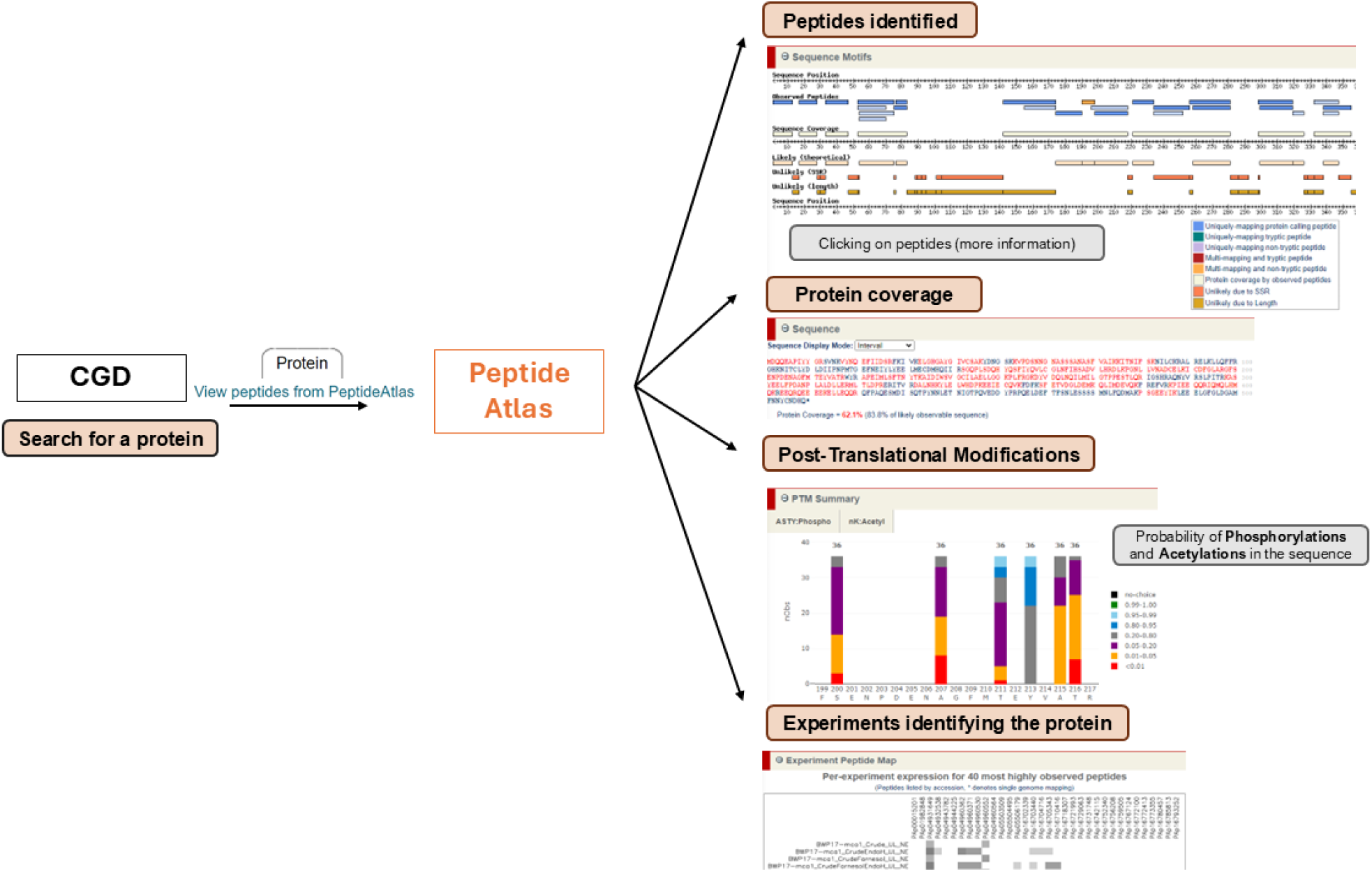
Schematic representation of the functionalities of the PeptideAtlas for a given *C. albicans* protein.

The main novelty beyond the expansion of the *Candida albicans* PeptideAtlas in this update is the ability to observe the probabilities of PTMs identified with information at an individual spectral level with hyperlinked access to the annotated spectra. Consequently, the focus has been placed on providing instructions on how to interpret and apply these results.

### c) Mapping biological PTMs: Phosphorylation and Acetylation

Several PXDs that had enrichment methods for physiologically important PTMs of phosphorylation and acetylation were selected (*Supplementary Table 1 (xlsx)*). A sophisticated PTM viewer in PeptideAtlas allows detailed examination of these PTMs, including direct links to all spectral matches. Here, we illustrate two examples of canonical proteins that undergo PTMs, being the substrate of phosphorylation and acetylation. Our summary in this publication is limited on PTMs to canonical proteins where high localization probability is appreciated, but all confidence levels of protein identification are available through the PeptideAtlas web interface. We recommend therefore to use the PeptideAtlas to evaluate specific PTM sites searched for if these are of particular interest for the reader.

*Figure 6* shows the functionality of PeptideAtlas for the determination of phospho-sites. Mitogen-activated protein kinase (Mkc1) is one example in which a protein was identified as a canonical protein and for which phosphorylation has important functional significance. The Mkc1p is part of the p42-44 MAP kinase family and is involved in the fungal cell integrity pathway, a signal transduction pathway known to be activated by cell wall stress. It has been demonstrated that Mkc1 is required for invasive hyphal growth and normal biofilm development. Its activation by phosphorylation occurs after membrane perturbation, cell wall stress and oxidative stress.^54,55^

**Figure 6.**
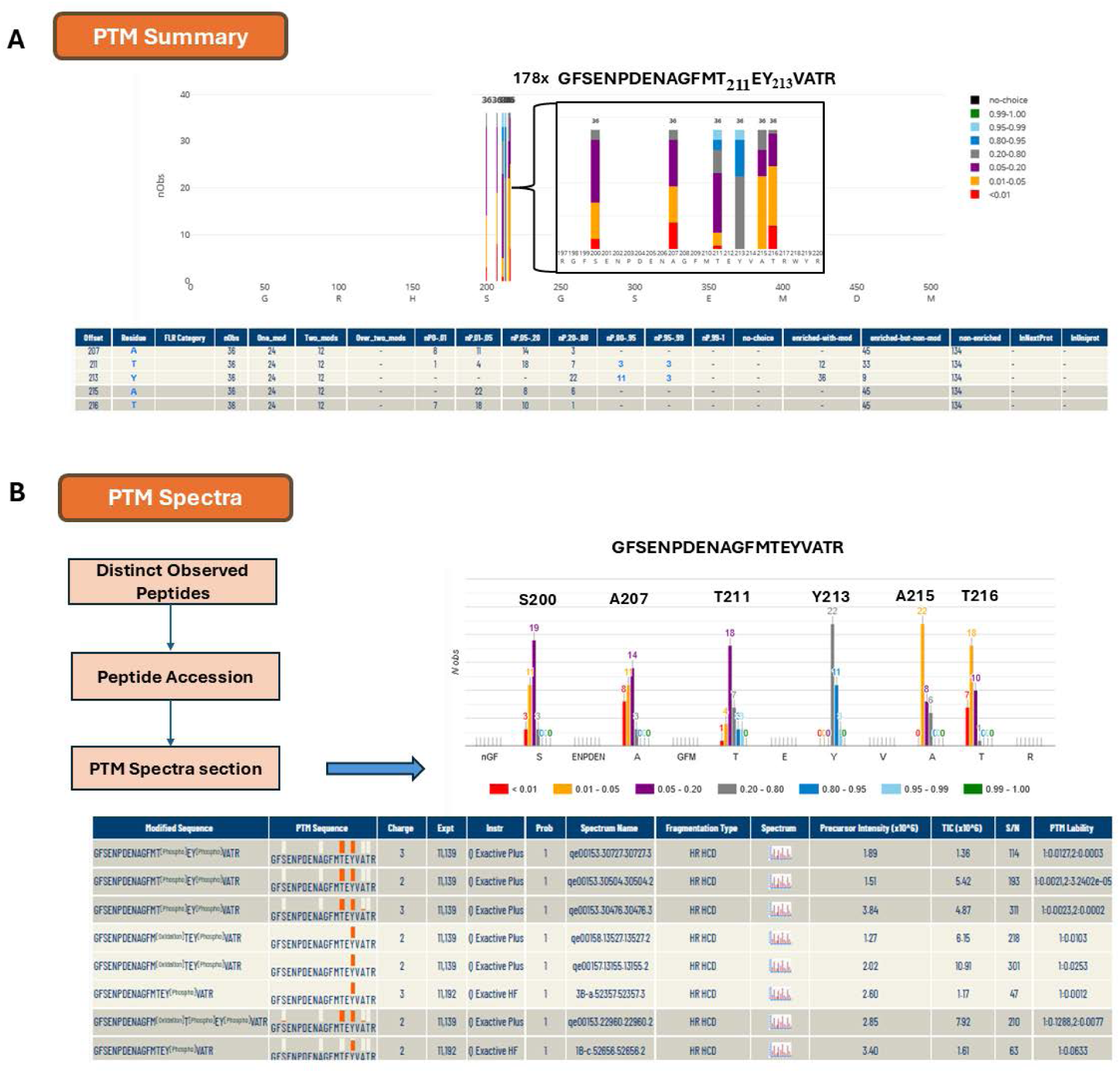
Illustration of the functionality of PeptideAtlas for the determination of phospho-sites based on the example of Mkc1 (CR_00120C_A). A) Probabilities of localization of phosphates on potential sites (as colored in bars) along the complete protein sequence. The phosphorylated residues in GFSENPDENAGFMTEYVATR peptide are highlighted with frequency of specific p-sites, color-coded by localization probability. This shows that the GFSENPDENAGFMTEYVATR peptide was observed 178 times and that T211 and Y213 were observed 36 total times each, 3 of them at the highest significance level. A numerical summary of p-site observations and information about specific sample enrichment for phospho-peptides based on metadata information collected from the individual PXD submissions is displayed below. Explanation for columns: Offset - residue offset in the protein; FLR Category, – FLR punctuation; Residue, - amino acid; nObs, - total observed PTM spectra for the site; One_mod, - the containing peptides have only one observed PTM site; Two_mods - the containing peptides have only two observed PTM site; Over_two_mods - the containing peptides have more than two observed PTM sites; nP0-.01 - number of observed PSM with PTMProphet probability < 0.01; nP.01-.05 - number of observed PSM with PTMProphet probability ≥ 0.01 and < 0.05; nP.05-.20 - number of observed PSM with PTMProphet probability ≥ 0.05 and < 0.2; nP.20-.80 - number of observed PSM with PTMProphet probability ≥ 0.2 and < 0.8; nP.80-.95 - number of observed PSM with PTMProphet probability ≥ 0.80 and < 0.95; nP.95-.99 - Number of observed PSM with PTMProphet probability ≥0.95 and < 0.99; nP.99-1 - PTMProphet probability ≥0.99; no choice - number of PSMs covering this site for which there was no choice in the localization of the PTM. Only one residue was an S, T, or Y; enriched-with-mod - number of PSMs covering this site with phospho modification on this site, and originating from a phospho-enriched sample; enriched-but-non-mod - number of PSMs covering this site with no phospho modification anywhere on the peptide anywhere on the peptide, but yet originating from a phospho-enriched sample; non-enriched - number of PSMs covering this site with no phospho modification anywhere on the peptide, and originating from a non-enriched sample, typically not searched for phospho; inNexProt - PTM site annotated in neXtprot; InUniProt - PTM site annotated in UniProt. B) Detailed view of the phosphorylated peptide GFSENPDENAGFMTEYVATR and phospho-site localization probability distributions. The lower panel shows information at an individual spectral level with hyperlinked access to the annotated spectra.

Mkc1 (CR_00120C_A) was identified with 34 distinct peptides and with a 62.1% protein sequence coverage. The *C. albicans* PeptideAtlas shows only one phosphorylated peptide: GFSENPDENAGFMTEYVATR identified 36 times. In 3 cases, both the phosphorylation sites at position T211 and Y213, were identified with a significance of 0.95 < P < 0.99 in light blue with additional observations for these sites at lower probabilities. These are serine S200 and threonine T216 whose phosphorylation sites were assigned with low probabilities, indicating with high confidence that the detected phosphate was not positioned at those sites.

In *Figure 6A* a numerical summary of the phosphorylation sites is displayed. Peptide GFSENPDENAGFMTEYVATR was detected 134 times in samples that were not enriched in phosphorylated peptides, 12 times as a single phosphorylated peptide and 24 times as a doubly phosphorylated peptide from enriched samples. T211 was identified exclusively in the doubly phosphorylated peptide. Y213 was identified in all 36 observations. *Figure 6B* shows a detailed peptide view of the phosphorylated peptide GFSENPDENAGFMTEYVATR and phospho-site identification scores. The lower panel in *Figure 6B* shows information at the individual spectral level with hyperlinked access to the annotated spectra. Together, this strongly suggests that Mkc1 is phosphorylated at T211 and Y213. Navarro-García et al^56^ established that phosphorylation of the threonine and tyrosine residues in the TEY signature of domain VIII of the p42-44 MAP kinase family is necessary for phosphorylation activity. When both residues are dephosphorylated, p42-44 MAP kinases are unable to phosphorylate their substrates.

*Figure 7* demonstrates the functionality of PeptideAtlas for the determination of acetylated residues. Hsp90 (C7_02030W_A) is one example in which a protein was identified as a canonical protein and for which acetylation was previously demonstrated to have important functional significance. Hsp90 is a conserved molecular chaperone that facilitates the folding and function of hundreds of proteins, many of which serve as core hubs of signal transduction networks. Hsp90 governs morphogenesis and virulence such that compromise of Hsp90 function induces the transition from yeast to filamentous growth in the absence of any additional inducing cue. Hsp90 governs temperature-dependent morphogenesis by inhibiting signaling through the cAMP-protein kinase A (PKA) pathway.^57^ Furthermore, it is necessary to highlight the important role of Hsp90 in mediating echinocandin and biofilm azole resistance.^58^

**Figure 7.**
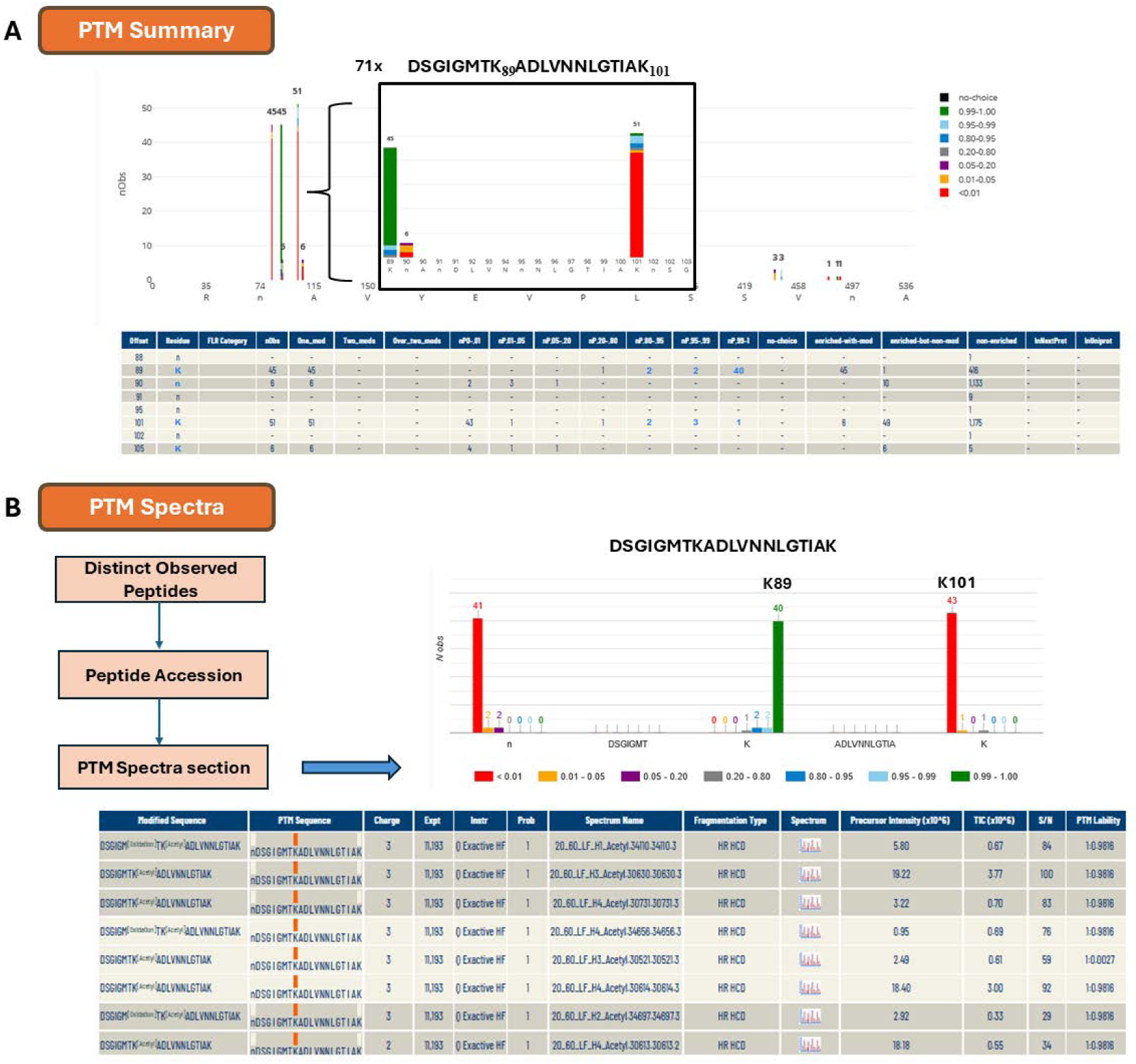
Illustration of the functionality of PeptideAtlas for the determination of acetylated residues based on the example of Hsp90 (C7_02030W_A). A) Probabilities of localization of acetylated sites in DSGIGMTKADLVNNLGTIAK peptide are highlighted color-coded by localization probability. This shows that the acetylated peptide was observed 71 times and that K89 and K101 were observed 45 and 51 total times respectively. A numerical summary of acetylated site observations and information about specific sample enrichment for acetylated peptides based on metadata information collected from each individual PXD submission is displayed below. Explanation for columns: Offset - residue offset in the protein; FLR Category: FLR punctuation; Residue - amino acid; nObs - total observed PTM spectra for the site; One_mod - the containing peptides have only one observed PTM site; Two_mods - the containing peptides have only two observed PTM site; Over_two_mods - the containing peptides have more than two observed PTM sites; nP0-.01 - number of observed PSM with PTMProphet probability < 0.01; nP.01-.05 - number of observed PSM with PTMProphet probability ≥ 0.01 and < 0.05; nP.05-.20 - number of observed PSM with PTMProphet probability ≥ 0.05 and < 0.2; nP.20-.80 - number of observed PSM with PTMProphet probability ≥ 0.2 and < 0.8; nP.80-.95 - number of observed PSM with PTMProphet probability ≥ 0.80 and < 0.95; nP.95-.99 - number of observed PSM with PTMProphet probability ≥0.95 and < 0.99; nP.99-1 - PTMProphet probability ≥0.99; no choice - Number of PSMs covering this site for which there was no choice in the localization of the PTM. Only one residue was an S, T, or Y; enriched-with-mod - number of PSMs covering this site with phospho modification on this site, and originating from a phospho-enriched sample; enriched-but-non-mod - number of PSMs covering this site with no acetyl modification anywhere on the peptide anywhere on the peptide, but yet originating from a acetyl-enriched sample; non-enriched - Number of PSMs covering this site with no acetyl modification anywhere on the peptide, and originating from a non-enriched sample, typically not searched for acetylation; inNexProt - PTM site annotated in neXtprot; InUniprot - PTM site annotated in UniProt. B) Detailed view of the acetylated peptide DSGIGMTKADLVNNLGTIAK and acetylation site localization probability distributions. The lower panel shows information at an individual spectral level with hyperlinked access to the annotated spectra.

Hsp90 (C7_02030W_A) was identified with 355 peptides and 96.1% protein sequence coverage. In *Figure 7* we present the acetylated peptide DSGIGMTKADLVNNLGTIAK identified 71 times. In 40 cases the acetylated K89 was identified with a significance of 0.99 < P < 1.00 with additional observations for this site at lower probability. There is another acetylation site at K101 assigned with low probability, indicating high confidence that the detected acetyl was not positioned at this site.

In *Figure 7A* a numerical summary of the acetylation sites is displayed. K89 was observed 416 times in samples that were not enriched for acetylated peptides and 45 times in a single acetylated peptide from enriched samples and 51 times for K101 in an acetylated peptide from enriched samples.

There are several peptides of C7_02030W_A where acetylation was observed, such as peptide LFLKEDQLEYLEEKR which was identified 1022 times. In 3 cases the acetylated K191 was identified with a significance of 0.99 < P < 1.00 with additional observations for this site at lower probability. The peptide VVVSYKLVDAPAAIR was identified 8 times, 3 of them the acetylated K567 was identified with a significance of 0.99 < P < 1.00 with additional observations for this site at lower probability.

## CONCLUSION

The 2024 update of the *C. albicans* PeptideAtlas (2024-03 version) that includes thirty-three new datasets chosen from PRIDE based on specific criteria (such as the use of SC5314 or mutants, LC-MS/MS, DDA mode, non-glycosylated enrichment, and CGD) to now include extensive identification of PTMs. Finally, these experiments were reanalyzed in conjunction with data from previous versions of PeptideAtlas using the TPP.

The 2024 the *C. albicans* PeptideAtlas build has resulted in a 12-fold increase in the number of identified PSMs, a 2.5-fold increase in characterized peptides, and a 1.2-fold increase in proteins compared to the previous build. A total of 11,925,423 PSM, 176,568 peptides and 4,955 canonical protein sequences make it the most comprehensive proteomics resource available up to date with a coverage of 80% of the total predicted proteome.

In addition, highly confident protein identifications have been made for 76% of the genes labeled as “uncharacterized” (without a known protein product) and 1% of the genes categorized as “dubious” (unlikely to encode a product) in CGD. A total of 3,382 phosphorylated serine and 536 phosphorylated threonine sites were identified, with false localization rates of 1% and 5.3%, and PTMProphet scores ranging between 0.95 and 0.99 respectively. The majority of spectra matched to peptides have a charge state of 2+ (57.19%) and the most common peptide length is 12 aa (7.36%).

Approximately 80% of the proteins belonging to nuclear chromosomes were identified as canonical proteins and 15% of them were not observed. An explanation of the PeptideAtlas tools and instructions on how to access them, either through the PeptideAtlas web interface or via the CGD, have been provided. Accessing PeptideAtlas via its own web interface offers information on the “Build overview”, “PTM coverage”, “Experiment contribution”, and “Dataset contribution”. Through a protein entry on the CGD webpage, users can view information about the peptides identified for that protein, protein coverage, probability of phosphorylation and acetylation within the sequence, and the experiments in which the protein has been identified. A detailed explanation on how to interpret and apply PTM detection and localization information was provided. This latest expanded *C. albicans* PeptideAtlas build provides a comprehensive compendium of results from community proteomics data that will enable further research on this important pathogen.

## AUTHOR INFORMATION

### Author Contributions

Leticia Gomez-Artiguez, Samuel de la Cámara-Fuentes and Zhi Sun contributed equally to this work.

### Notes

The authors declare no competing financial interest.

## ASSOCIATED CONTENT

### Data Availability Statement

The mass spectrometry proteomics data used for the PeptideAtlas update is deposited at the ProteomeXchange Consortium via the PRIDE partner repository whose data set identifiers are listed in *Supplementary Table 1 (xlsx)*.

### Supporting Information

Supplementary Table 1: Summary information on the 33 selected PXD datasets for the PeptideAtlas update (.xlsx)

Supplementary Figure S1: Accumulative PXDs (PRIDE) with verified *C. albicans* content by year (2015-03/2024-04); Supplementary Table 2: Technical definition of protein identification confidence categories in the *C. albicans* PeptideAtlas build (.docx)

## ACKNOWLEDGMENT

This work was granted by PID2021-124062NB-I00 (CG) funded by Spanish Ministry of Science and Innovation/ State Research Agency 10.13039/501100011033, the National Institutes of Health grants R01 GM087221 (EWD, RLM), U19 AG023122 (RLM), S10 OD026936 (RLM) and by the National Science Foundation grants DBI-1933311 (EWD) and MRI-1920268 (RLM).

L. Gomez-Artiguez was supported by a contract funded by grant CT19/23-INVM-70 from the European Union-NextGenerationEU, as part of Investigo program in the frame of “Recovery, Transformation and Resilience Plan” co-funded by the Ministry of Labor and Social Economy and the National Public Employment Service. S. de la Cámara-Fuentes was supported by PTA2022-022414-I from the Spanish Ministry of Science and Innovation and the State Research Agency, along with REACT-UE ANTICIPA-CM from the Community of Madrid.

## ABBREVIATIONS

CGD: Candida Genome Database
DDA: Data-dependent analysis
FDR: False discovery rate
LC-MS/MS: liquid chromatography tandem mass spectrometry
PRIDE: Proteomics identification database
PSM: Peptide spectrum match
PTM: Post-translational modifications
TMT: Tandem mass tag
TPP: Trans-Proteomic pipeline

## REFERENCES

(1) Tsui, C.; Kong, E. F.; Jabra-Rizk, M. A. Pathogenesis of *Candida Albicans* Biofilm. Pathog. Dis. 2016, 74 (4), ftw018. 10.1093/femspd/ftw018.

(2) Talapko, J.; Juzbašić, M.; Matijević, T.; Pustijanac, E.; Bekić, S.; Kotris, I.; Škrlec, I. *Candida albicans*—The Virulence Factors and Clinical Manifestations of Infection. J Fungi (Basel). 2021 Jan 22;7(2):79. doi: 10.3390/jof7020079.

(3) Zhang, Z.; Zhu, R.; Luan, Z.; Ma, X. Risk of Invasive Candidiasis with Prolonged Duration of ICU Stay: A Systematic Review and Meta-Analysis. BMJ Open 2020, 10 (7), e036452. 10.1136/bmjopen-2019-036452.

(4) Hui, Y. Z.; Gai, S. Y.; Yu, L. R. A Ten-Year Retrospective Study of Invasive Candidiasis in a Tertiary Hospital in Beijing. Biomed Env. Sci. 2021 Oct 20;34(10):773–788. doi: 10.3967/bes2021.107.

(5) Parums, D. V. Editorial: The World Health Organization (WHO) Fungal Priority Pathogens List in Response to Emerging Fungal Pathogens During the COVID-19 Pandemic. Med. Sci. Monit. 2022, 28. 10.12659/MSM.939088.

(6) Vialas, V.; Sun, Z.; Loureiro Y Penha, C. V.; Carrascal, M.; Abián, J.; Monteoliva, L.; Deutsch, E. W.; Aebersold, R.; Moritz, R. L.; Gil, C. A Candida albicans PeptideAtlas. J. Proteomics 2014, 97, 62–68. 10.1016/j.jprot.2013.06.020.

(7) Deutsch EW, Bandeira N, Perez-Riverol Y, Sharma V, Carver JJ, Mendoza L, Kundu DJ, Wang S, Bandla C, Kamatchinathan S, Hewapathirana S, Pullman BS, Wertz J, Sun Z, Kawano S, Okuda S, Watanabe Y, MacLean B, MacCoss MJ, Zhu Y, Ishihama Y, Vizcaíno JA. The ProteomeXchange consortium at 10 years: 2023 update. Nucleic Acids Res. 2023 Jan 6;51(D1):D1539–D1548. doi: 10.1093/nar/gkac1040

(8) Deutsch, E. W.; Mendoza, L.; Shteynberg, D.; Slagel, J.; Sun, Z.; Moritz, R. L. Trans-Proteomic Pipeline, a Standardized Data Processing Pipeline for Large-Scale Reproducible Proteomics Informatics. Proteomics Clin Appl. 2015 Aug;9(7-8):745–54. doi: 10.1002/prca.201400164.

(9) Desiere, F.; Deutsch, E. W.; Nesvizhskii, A. I.; Mallick, P.; King, N. L.; Eng, J. K.; Aderem, A.; Boyle, R.; Brunner, E.; Donohoe, S.; Fausto, N.; Hafen, E.; Hood, L.; Katze, M. G.; Kennedy, K. A.; Kregenow, F.; Lee, H.; Lin, B.; Martin, D.; Ranish, J. A.; Rawlings, D. J.; Samelson, L. E.; Shiio, Y.; Watts, J. D.; Wollscheid, B.; Wright, M. E.; Yan, W.; Yang, L.; Yi, E. C.; Zhang, H.; Aebersold, R. Integration with the Human Genome of Peptide Sequences Obtained by High-Throughput Mass Spectrometry. Genome Biol. 2005;6(1):R9. doi: 10.1186/gb-2004-6-1-r9.

(10) Hesselager, M. O.; Codrea, M. C.; Sun, Z.; Deutsch, E. W.; Bennike, T. B.; Stensballe, A.; Bundgaard, L.; Moritz, R. L.; Bendixen, E. The Pig PeptideAtlas: A Resource for Systems Biology in Animal Production and Biomedicine. Proteomics. 2016 Feb;16(4):634–44. doi: 10.1002/pmic.201500195.

(11) McCord, J.; Sun, Z.; Deutsch, E. W.; Moritz, R. L.; Muddiman, D. C. The PeptideAtlas of the Domestic Laying Hen. J Proteome Res. 2017 Mar 3;16(3):1352–1363. doi: 10.1021/acs.jproteome.6b00952.

(12) Van Wijk, K. J.; Leppert, T.; Sun, Q.; Boguraev, S. S.; Sun, Z.; Mendoza, L.; Deutsch, E. W. The Arabidopsis PeptideAtlas: Harnessing Worldwide Proteomics Data to Create a Comprehensive Community Proteomics Resource. Plant Cell 2021, 33 (11), 3421–3453. 10.1093/plcell/koab211.

(13) Van Wijk KJ, Leppert T, Sun Z, Guzchenko I, Debley E, Sauermann G, Routray P, Mendoza L, Sun Q, Deutsch EW. The *Zea mays* PeptideAtlas: A New Maize Community Resource. J Proteome Res. 2024 Sep 6;23(9):3984–4004. doi: 10.1021/acs.jproteome.4c00320.

(14) King, N. L.; Deutsch, E. W.; Ranish, J. A.; Nesvizhskii, A. I.; Eddes, J. S.; Mallick, P.; Eng, J.; Desiere, F.; Flory, M.; Martin, D. B.; Kim, B.; Lee, H.; Raught, B.; Aebersold, R. Analysis of the Saccharomyces Cerevisiae Proteome with PeptideAtlas. Genome Biol. 2006; 7(11):R106. doi: 10.1186/gb-2006-7-11-r106.

(15) Gunaratne J, Schmidt A, Quandt A, Neo SP, Saraç OS, Gracia T, Loguercio S, Ahrné E, Xia RL, Tan KH, Lössner C, Bähler J, Beyer A, Blackstock W, Aebersold R. Extensive mass spectrometry-based analysis of the fission yeast proteome: the Schizosaccharomyces pombe PeptideAtlas. Mol Cell Proteomics. 2013 Jun;12(6):1741–51. doi: 10.1074/mcp.M112.023754.

(16) Michalik S, Depke M, Murr A, Gesell Salazar M, Kusebauch U, Sun Z, Meyer TC, Surmann K, Pförtner H, Hildebrandt P, Weiss S, Palma Medina LM, Gutjahr M, Hammer E, Becher D, Pribyl T, Hammerschmidt S, Deutsch EW, Bader SL, Hecker M, Moritz RL, Mäder U, Völker U, Schmidt F. A global Staphylococcus aureus proteome resource applied to the in vivo characterization of host-pathogen interactions. Sci Rep. 2017 Sep 8;7(1):9718. doi: 10.1038/s41598-017-10059-w.

(17) Reales-Calderón JA, Sun Z, Mascaraque V, Pérez-Navarro E, Vialás V, Deutsch EW, Moritz RL, Gil C, Martínez JL, Molero G. A wide-ranging Pseudomonas aeruginosa PeptideAtlas build: A useful proteomic resource for a versatile pathogen. J Proteomics. 2021 May 15;239:104192. doi: 10.1016/j.jprot.2021.104192.

(18) Vialas, V.; Sun, Z.; Reales-Calderón, J. A.; Hernáez, M. L.; Casas, V.; Carrascal, M.; Abián, J.; Monteoliva, L.; Deutsch, E. W.; Gil, C. A Comprehensive *Candida albicans* PeptideAtlas Build Enables Deep Proteome Coverage. J Proteomics 2016 Jan 10;131:122–130. doi: 10.1016/j.jprot.2015.10.019.

(19) Perez-Riverol, Y.; Bai, J.; Bandla, C.; García-Seisdedos, D.; Hewapathirana, S.; Kamatchinathan, S.; Kundu, D. J.; Prakash, A.; Frericks-Zipper, A.; Eisenacher, M.; Walzer, M.; Wang, S.; Brazma, A.; Vizcaíno, J. A. The PRIDE Database Resources in 2022: A Hub for Mass Spectrometry-Based Proteomics Evidences. Nucleic Acids Res. 2022, 50 (D1), D543–D552. 10.1093/nar/gkab1038.

(20) Keller, A.; Eng, J.; Zhang, N.; Li, X.; Aebersold, R. A Uniform Proteomics MS/MS Analysis Platform Utilizing Open XML FIle Formats. 2005; 1:2005.0017. doi: 10.1038/msb4100024.

(21) Hulstaert, N.; Shofstahl, J.; Sachsenberg, T.; Walzer, M.; Barsnes, H.; Martens, L.; Perez-Riverol, Y. ThermoRawFileParser: Modular, Scalable and Cross-Platform RAW File Conversion. J Proteome Res. 2020 Jan 3;19(1):537–542. doi: 10.1021/acs.jproteome.9b00328.

(22) Kong, A. T.; Leprevost, F. V.; Avtonomov, D. M.; Mellacheruvu, D.; Nesvizhskii, A. I. MSFragger: Ultrafast and Comprehensive Peptide Identification in Shotgun Proteomics. Nat Methods 2017. May;14(5):513–520. doi: 10.1038/nmeth.4256.

(23) UniProt Consortium. UniProt: A Worldwide Hub of Protein Knowledge. Nucleic Acids Res. 2019, 47 (D1), D506–D515. 10.1093/nar/gky1049.

(24) Slagel, J.; Mendoza, L.; Shteynberg, D.; Deutsch, E. W.; Moritz, R. L. Processing Shotgun Proteomics Data on the Amazon Cloud with the Trans-Proteomic Pipeline. Mol. Cell. Proteomics 2015, 14 (2), 399–404. 10.1074/mcp.O114.043380.

(25) Ramsbottom, K. A.; Prakash, A.; Riverol, Y. P.; Camacho, O. M.; Martin, M.-J.; Vizcaíno, J. A.; Deutsch, E. W.; Jones, A. R. Method for Independent Estimation of the False Localization Rate for Phosphoproteomics. J Proteome Res. 2022 Jul 1;21(7):1603–1615. doi: 10.1021/acs.jproteome.1c00827.

(26) Keller, A.; Nesvizhskii, A. I.; Kolker, E.; Aebersold, R. Empirical Statistical Model To Estimate the Accuracy of Peptide Identifications Made by MS/MS and Database Search. Anal. Chem. 2002, 74 (20), 5383–5392. 10.1021/ac025747h.

(27) Shteynberg, D.; Deutsch, E. W.; Lam, H.; Eng, J. K.; Sun, Z.; Tasman, N.; Mendoza, L.; Moritz, R. L.; Aebersold, R.; Nesvizhskii, A. I. iProphet: Multi-Level Integrative Analysis of Shotgun Proteomic Data Improves Peptide and Protein Identification Rates and Error Estimates. Mol. Cell. Proteomics 2011, 10 (12), M111.007690. 10.1074/mcp.M111.007690.

(28) Shteynberg, D. D.; Deutsch, E. W.; Campbell, D. S.; Hoopmann, M. R.; Kusebauch, U.; Lee, D.; Mendoza, L.; Midha, M. K.; Sun, Z.; Whetton, A. D.; Moritz, R. L. PTMProphet: Fast and Accurate Mass Modification Localization for the Trans-Proteomic Pipeline. J. Proteome Res. 2019, 18 (12), 4262–4272. 10.1021/acs.jproteome.9b00205.

(29) Reiter, L.; Claassen, M.; Schrimpf, S. P.; Jovanovic, M.; Schmidt, A.; Buhmann, J. M.; Hengartner, M. O.; Aebersold, R. Protein Identification False Discovery Rates for Very Large Proteomics Data Sets Generated by Tandem Mass Spectrometry. Mol. Cell. Proteomics 2009, 8 (11), 2405–2417. 10.1074/mcp.M900317-MCP200.

(30) Farrah, T.; Deutsch, E. W.; Omenn, G. S.; Campbell, D. S.; Sun, Z.; Bletz, J. A.; Mallick, P.; Katz, J. E.; Malmström, J.; Ossola, R.; Watts, J. D.; Lin, B.; Zhang, H.; Moritz, R. L.; Aebersold, R. A High-Confidence Human Plasma Proteome Reference Set with Estimated Concentrations in PeptideAtlas. Mol. Cell. Proteomics MCP 2011, 10 (9), M110.006353. 10.1074/mcp.M110.006353.

(31) Deutsch, E. W.; Lane, L.; Overall, C. M.; Bandeira, N.; Baker, M. S.; Pineau, C.; Moritz, R. L.; Corrales, F.; Orchard, S.; Van Eyk, J. E.; Paik, Y.-K.; Weintraub, S. T.; Vandenbrouck, Y.; Omenn, G. S. Human Proteome Project Mass Spectrometry Data Interpretation Guidelines 3.0. J. Proteome Res. 2019, 18 (12), 4108–4116. 10.1021/acs.jproteome.9b00542.

(32) Willger, S. D.; Liu, Z.; Olarte, R. A.; Adamo, M. E.; Stajich, J. E.; Myers, L. C.; Kettenbach, A N.; Hogan, D. A. Analysis of the *Candida albicans* Phosphoproteome. Eukaryot. Cell 2015, 14 (5), 474–485. 10.1128/EC.00011-15.

(33) Luo, T.; Krüger, T.; Knüpfer, U.; Kasper, L.; Wielsch, N.; Hube, B.; Kortgen, A.; Bauer, M.; Giamarellos-Bourboulis, E. J.; Dimopoulos, G.; Brakhage, A. A.; Kniemeyer, O. Immunoproteomic Analysis of Antibody Responses to Extracellular Proteins of *Candida albicans* Revealing the Importance of Glycosylation for Antigen Recognition. J. Proteome Res. 2016, 15 (8), 2394–2406. 10.1021/acs.jproteome.5b01065.

(34) Léger, T.; Garcia, C.; Camadro, J.-M. The Metacaspase (Mca1p) Restricts O-Glycosylation During Farnesol-Induced Apoptosis in *Candida albicans*. Mol. Cell. Proteomics 2016, 15 (7), 2308–2323. 10.1074/mcp.M116.059378.

(35) Leach, M. D.; Kim, T.; DiGregorio, S. E.; Collins, C.; Zhang, Z.; Duennwald, M. L.; Cowen, L. E. *Candida Albicans* Is Resistant to Polyglutamine Aggregation and Toxicity. G3 GenesGenomesGenetics 2017, 7 (1), 95–108. 10.1534/g3.116.035675.

(36) She, X.; Zhang, P.; Gao, Y.; Zhang, L.; Wang, Q.; Chen, H.; Calderone, R.; Liu, W.; Li, D. A Mitochondrial Proteomics View of Complex I Deficiency in *Candida albicans*. Mitochondrion 2018, 38, 48–57. 10.1016/j.mito.2017.08.003.

(37) Munusamy, K.; Loke, M. F.; Vadivelu, J.; Tay, S. T. LC-MS Analysis Reveals Biological and Metabolic Processes Essential for *Candida albicans* Biofilm Growth. Microb. Pathog. 2021, 152, 104614. 10.1016/j.micpath.2020.104614.

(38) Duval, C.; Macabiou, C.; Garcia, C.; Lesuisse, E.; Camadro, J.; Auchère, F. The Adaptive Response to Iron Involves Changes in Energetic Strategies in the Pathogen *Candida Albicans*. MicrobiologyOpen 2020, 9 (2), e970. 10.1002/mbo3.970.

(39) Arita, G. S.; Meneguello, J. E.; Sakita, K. M.; Faria, D. R.; Pilau, E. J.; Ghiraldi-Lopes, L. D.; Campanerut-Sá, P. A. Z.; Kioshima, É. S.; Bonfim-Mendonça, P. D. S.; Svidzinski, T. I. E. Serial Systemic *Candida albicans* Infection Highlighted by Proteomics. Front. Cell. Infect. Microbiol. 2019, 9, 230. 10.3389/fcimb.2019.00230.

(40) Vaz, C.; Pitarch, A.; Gómez-Molero, E.; Amador-García, A.; Weig, M.; Bader, O.; Monteoliva, L.; Gil, C. Mass Spectrometry-Based Proteomic and Immunoproteomic Analyses of the *Candida albicans* Hyphal Secretome Reveal Diagnostic Biomarker Candidates for Invasive Candidiasis. J. Fungi 2021, 7 (7), 501. 10.3390/jof7070501.

(41) Balachandra, V. K.; Verma, J.; Shankar, M.; Tucey, T. M.; Traven, A.; Schittenhelm, R. B.; Ghosh, S. K. The RSC (Remodels the Structure of Chromatin) Complex of *Candida albicans* Shows Compositional Divergence with Distinct Roles in Regulating Pathogenic Traits. PLOS Genet. 2020, 16 (11), e1009071. 10.1371/journal.pgen.1009071.

(42) Childers, D. S.; Avelar, G. M.; Bain, J. M.; Pradhan, A.; Larcombe, D. E.; Netea, M. G.; Erwig, L. P.; Gow, N. A. R.; Brown, A. J. P. Epitope Shaving Promotes Fungal Immune Evasion. mBio 2020, 11 (4), e00984–20. 10.1128/mBio.00984-20.

(43) Jenull, S.; Mair, T.; Tscherner, M.; Penninger, P.; Zwolanek, F.; Silao, F.-G. S.; De San Vicente, K. M.; Riedelberger, M.; Bandari, N. C.; Shivarathri, R.; Petryshyn, A.; Chauhan, N.; Zacchi, L. F.; -Landmann, S. L.; Ljungdahl, P. O.; Kuchler, K. The Histone Chaperone HIR Maintains Chromatin States to Control Nitrogen Assimilation and Fungal Virulence. Cell Rep. 2021, 36 (3), 109406. 10.1016/j.celrep.2021.109406.

(44) Zhou, X.; Song, N.; Li, D.; Li, X.; Liu, W. Systematic Analysis of the Lysine Crotonylome and Multiple Posttranslational Modification Analysis (Acetylation, Succinylation, and Crotonylation) in Candida albicans. mSystems 2021, 6 (1), e01316–20. 10.1128/mSystems.01316-20.

(45) Amador-García, A.; Zapico, I.; Borrajo, A.; Malmström, J.; Monteoliva, L.; Gil, C. Extending the Proteomic Characterization of *Candida albicans* Exposed to Stress and Apoptotic Inducers through Data-Independent Acquisition Mass Spectrometry. mSystems 2021, 6 (5), e00946–21. 10.1128/mSystems.00946-21.

(46) Martínez-López, R.; Hernáez, M. L.; Redondo, E.; Calvo, G.; Radau, S.; Pardo, M.; Gil, C.; Monteoliva, L. *Candida albicans* Hyphal Extracellular Vesicles Are Different from Yeast Ones, Carrying an Active Proteasome Complex and Showing a Different Role in Host Immune Response. Microbiol. Spectr. 2022, 10 (3), e00698–22. 10.1128/spectrum.00698-22.

(47) Zheng, H.; Song, N.; Zhou, X.; Mei, H.; Li, D.; Li, X.; Liu, W. Proteome-Wide Analysis of Lysine 2-Hydroxyisobutyrylation in *Candida albicans*. mSystems 2021, 6 (1), e01129–20. 10.1128/mSystems.01129-20.

(48) Zhang, Y.; Tang, C.; Zhang, Z.; Li, S.; Zhao, Y.; Weng, L.; Zhang, H. Deletion of the ATP2 Gene in *Candida albicans* Blocks Its Escape From Macrophage Clearance. Front. Cell. Infect. Microbiol. 2021, 11, 643121. 10.3389/fcimb.2021.643121.

(49) Łabędzka-Dmoch, K.; Kolondra, A.; Karpińska, M. A.; Dębek, S.; Grochowska, J.; Grochowski, M.; Piątkowski, J.; Hoang Diu Bui, T.; Golik, P. Pervasive Transcription of the Mitochondrial Genome in *Candida albicans* Is Revealed in Mutants Lacking the mtEXO RNase Complex. RNA Biol. 2021, 18 (sup1), 303–317. 10.1080/15476286.2021.1943929.

(50) Böttcher, B.; Driesch, D.; Krüger, T.; Garbe, E.; Gerwien, F.; Kniemeyer, O.; Brakhage, A. A.; Vylkova, S. Impaired Amino Acid Uptake Leads to Global Metabolic Imbalance of *Candida albicans* Biofilms. Npj Biofilms Microbiomes 2022, 8 (1), 78. 10.1038/s41522-022-00341-9.

(51) Brandt, P.; Gerwien, F.; Wagner, L.; Krüger, T.; Ramírez-Zavala, B.; Mirhakkak, M. H.; Schäuble, S.; Kniemeyer, O.; Panagiotou, G.; Brakhage, A. A.; Morschhäuser, J.; Vylkova, S. *Candida albicans* SR-Like Protein Kinases Regulate Different Cellular Processes: Sky1 Is Involved in Control of Ion Homeostasis, While Sky2 Is Important for Dipeptide Utilization. Front. Cell. Infect. Microbiol. 2022, 12, 850531. 10.3389/fcimb.2022.850531.

(52) Zarnowski, R.; Noll, A.; Chevrette, M. G.; Sanchez, H.; Jones, R.; Anhalt, H.; Fossen, J.; Jaromin, A.; Currie, C.; Nett, J. E.; Mitchell, A.; Andes, D. R. Coordination of Fungal Biofilm Development by Extracellular Vesicle Cargo. Nat. Commun. 2021, 12 (1), 6235. 10.1038/s41467-021-26525-z.

(53) Li, S.; Zhao, Y.; Zhang, Y.; Zhang, Y.; Zhang, Z.; Tang, C.; Weng, L.; Chen, X.; Zhang, G.; Zhang, H. The δ Subunit of F1Fo-ATP Synthase Is Required for Pathogenicity of *Candida albicans*. Nat. Commun. 2021, 12 (1), 6041. 10.1038/s41467-021-26313-9.

(54) Kumamoto, C. A. A Contact-Activated Kinase Signals *Candida Albicans* Invasive Growth and Biofilm Development. Proc. Natl. Acad. Sci. 2005, 102 (15), 5576–5581. 10.1073/pnas.0407097102.

(55) Navarro-Garcia, F.; Eisman, B.; Fiuza, S. M. The MAP Kinase Mkc1p Is Activated under Different Stress Conditions in Candida albicans. 2005.

(56) Navarro-García, F.; Sánchez, M.; Pla, J.; Nombela, C. Functional Characterization of the *MKC1* Gene of *Candida Albicans*, Which Encodes a Mitogen-Activated Protein Kinase Homolog Related to Cell Integrity. Mol. Cell. Biol. 1995, 15 (4), 2197–2206. 10.1128/MCB.15.4.2197.

(57) Robbins, N.; Cowen, L. E. Roles of Hsp90 in *Candida albicans* Morphogenesis and Virulence. Curr. Opin. Microbiol. 2023, 75, 102351. 10.1016/j.mib.2023.102351.

(58) Singh, S. D.; Robbins, N.; Zaas, A. K.; Schell, W. A.; Perfect, J. R.; Cowen, L. E. Hsp90 Governs Echinocandin Resistance in the Pathogenic Yeast *Candida albicans* via Calcineurin. PLoS Pathog. 2009, 5 (7), e1000532. 10.1371/journal.ppat.1000532.

