## Supplementary Figure 1 and Supplementary Table 2 for "*Candida albicans*: a comprehensive view of the proteome"

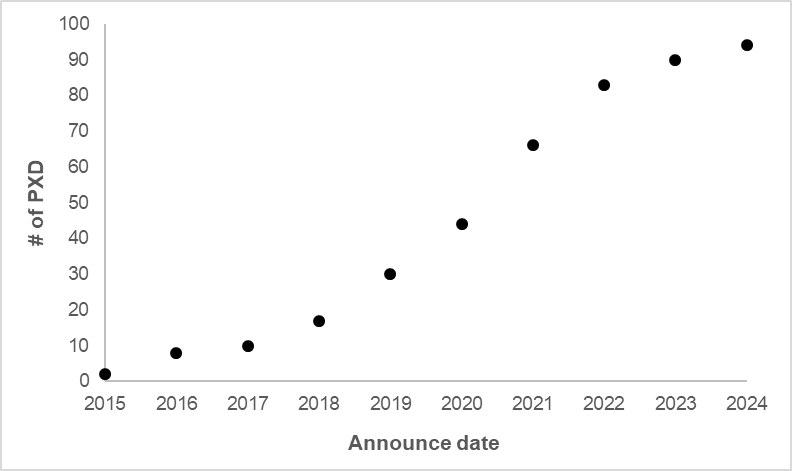

**Supplementary Figure 1.** Accumulative PXDs (PRIDE) with verified *C. albicans* content by year (2015-03/2024-04).

**Supplementary Table 2**. Technical definition of protein identification confidence categories in the *C. albicans* PeptideAtlas build.

| **Protein label** | **Technical definition** |
| --- | --- |
| Canonical | Proteins with at least two 9AA or greater peptides with a total extent of 18AA or greater that are uniquely mapping within the core reference proteome. |
| Noncore-Canonical | Proteins with at least 9AA or greater peptides with a total extent of 18AA or greater that do not map in the core reference proteome, but rather to an isoform, contaminant, or other protein missing from the core reference proteome. |
| Indistinguishable Representative | Protein has no unique peptides, and there are several indistinguishable proteins. The former is assigned to be an Indistinguishable Representative, while the latter are Indistinguishable |
| Insufficient Evidence | Protein has more unique peptides than shared peptides, but none are 9AA or greater. |
| Marginally Distinguished | Protein has unique peptides, but there are not more unique peptides than shared peptides, and the extended length of unique peptide is < 18AA. |
| Weak | Protein has more unique peptides than shared peptides, and only one uniquely mapping peptide 9AA or greater. |
| Not observed | Protein has no peptides above our PSM significance threshold. It may have PSMs of low significance, but these are not considered. |
